# Plant respiration: controlled by photosynthesis or biomass?

**DOI:** 10.1101/705400

**Authors:** Alessio Collalti, Mark G. Tjoelker, Günter Hoch, Annikki Mäkelä, Gabriele Guidolotti, Mary Heskel, Giai Petit, Michael G. Ryan, Giovanna Battipaglia, I. Colin Prentice

**Affiliations:** Institute for Agriculture and Forestry Systems in the Mediterranean, National Research Council of Italy (CNR-ISAFOM), 87036, Rende (CS), Italy; Department of Innovation in Biological, Agro-food and Forest Systems, University of Tuscia, 01100 Viterbo, Italy; Hawkesbury Institute for the Environment, Western Sydney University, Penrith, NSW 2751, Australia.; Department of Environmental Sciences – Botany, University of Basel, Schӧnbeinstrasse 6, Basel 4056, Switzerland.; Institute for Atmospheric and Earth System Research (INAR), Faculty of Science & and Faculty of Agriculture and Forestry, P.O. Box 27 (Latokartanonkaari 7) FI-00014 University of Helsinki, Finland.; Institute of Research on Terrestrial Ecosystem, National Research Council of Italy (CNR-IRET), 00015, Monterotondo Scalo (RM), Italy.; Department of Biology, Macalester College, Saint Paul, MN USA 55105.; Department of Land, Environment, Agriculture and Forestry, University of Padova, 35020 Legnaro (PD), Italy.; Natural Resource Ecology Laboratory, Colorado State University, Fort Collins, CO 80523-1499, USA.; USDA Forest Service, Rocky Mountain Experiment Station, Fort Collins, CO 80526, USA; Department of Environmental, Biological and Pharmaceutical Sciences and Technologies, University of Campania «L. Vanvitelli», 81100, Caserta, Italy.; AXA Chair of Biosphere and Climate Impacts, Imperial College London, Department of Life Sciences, Silwood Park Campus, Buckhurst Road, Ascot SL5 7PY, UK; Department of Biological Sciences, Macquarie University, North Ryde, NSW 2109, Australia; Ministry of Education Key Laboratory for Earth System Modeling, Department of Earth System Science, Tsinghua University, Beijing 100084, China

**Keywords:** Plant respiration, biomass accumulation, carbon use efficiency, gross primary production, net primary production, maintenance respiration, non-structural carbohydrates, metabolic scaling theory.

## Abstract

Two simplifying hypotheses have been proposed for whole-plant respiration. One links respiration to photosynthesis; the other to biomass. Using a first-principles carbon balance model with a prescribed live woody biomass turnover, applied at a forest research site where multidecadal measurements are available for comparison, we show that if turnover is fast the accumulation of respiring biomass is low and respiration depends primarily on photosynthesis; while if turnover is slow the accumulation of respiring biomass is high and respiration depends primarily on biomass. But the first scenario is inconsistent with evidence for substantial carryover of fixed carbon between years, while the second implies far too great an increase in respiration during stand development – leading to depleted carbohydrate reserves and an unrealistically high mortality risk. These two mutually incompatible hypotheses are thus both incorrect. Respiration is *not* linearly related either to photosynthesis or to biomass, but it is more strongly controlled by recent photosynthates (and reserve availability) than by total biomass.

## Introduction

The amount of carbon that accumulates in actively growing stands of vegetation depends on the balance of photosynthesis (gross primary production, *P*) and whole-plant (autotrophic) respiration (*R*). The difference between these fluxes is net primary production (*P*_n_). Most annual *P*_n_ is allocated to structural growth (*G*), but some is stored as non-structural carbohydrates (NSC, mostly starch and sugars), some is released back to the atmosphere in the form of biogenic volatile organic compounds (BVOCs), and some is exuded to the rhizosphere (Chapin *et al.* 2006). The fraction of *P* that accumulates in biomass, and the fraction that returns to the atmosphere through plant metabolism, are crucial quantities that determine the sign and magnitude of the global climate-carbon feedback – which remains one of the greatest sources of uncertainty in the global carbon cycle (Friedlingstein *et al.* 2014). But despite many ecophysiological studies aiming to understand *P*_n_ and *R* dynamics during stand development, a general understanding is still lacking.

Some authors have hypothesized a constant *P*_n_*:P* (carbon use efficiency, equivalent to 1 – (*R*:*P*)) ratio, with *R* tightly constrained by *P* irrespective of biomass, climate, tree species and stand age (e.g. Gifford 2003; Van Oijen *et al.* 2010). Waring *et al*. (1998, W98 hereafter) indicated a universal *P*_n_*:P* of ∼ 0.5. Since, ultimately, *R* depends on the matter produced by photosynthesis, Gifford (2003) suggested that these two processes must be tightly balanced over the longer term – making *R* proportional to *P*, consistent with W98. He argued that prescribing *P*_n_ (or *R*) as a constant fraction of *P* could be a simpler, and potentially more accurate, alternative to explicit, process-based modelling of *R*. A number of land vegetation models (reviewed in Collalti *&* Prentice 2019) adopt this simplification.

An alternative hypothesis, grounded in metabolic scaling theory, suggests that *R* should scale with biomass following a power law, (West *et al*. 1999). According to some studies (e.g. Reich *et al*. 2006, R06 hereafter), *R* (*Y*) scales isometrically (*b* ∼ 1) with whole-plant carbon (C) or nitrogen (N) contents (*X*), and this scaling is similar within and among different species, and irrespective of environmental and climatic conditions – which might influence the normalization constant (*a*), but not the exponent (*b*). Isometric scaling of *R* with biomass was assumed in the traditional view of forest dynamics set out e.g. by Kira *&* Shidei (1967) and Odum (1969). In the absence of major disturbances, if *R* increases in parallel with biomass, then *P*_n_ necessarily declines – because ultimately *P* cannot increase indefinitely, but rather stabilizes at canopy closure. Mori *et al*. (2010) however indicated that biomass and *R* are isometrically related only in young trees, tending towards *b* ∼ ¾ in mature trees. A general value of ⅗ has also been proposed (Michaletz *et al*. 2014). But however it is interpreted, this scaling hypothesis implies that *R* depends on biomass, and is related to *P* only to the extent that *P* and biomass vary together.

Although many terrestrial vegetation models simulate plant respiration assuming *R* to be a fixed fraction of *P*, others more explicitly couple *R* to biomass and thus only indirectly to *P*. The most widely used (and observationally supported) mechanistic approach, also adopted here, divides *R* into growth (*R*_G_) and maintenance (*R*_M_) components (McCree 1970; Thornley 1970). *R*_G_ is considered to be a fixed fraction of new tissue growth, independent of temperature, the fraction varying only with the cost of building the compounds constituting the new tissue (Penning de Vries 1972). Temperature, substrate availability and the demand for respiratory products are considered to control *R*_M_ (Cannell & Thornley 2000). Several studies have investigated the effects of short- and long-term changes in temperature on *R*_M_, mostly at the leaf level (e.g. Heskel *et al.* 2016; Huntingford *et al.* 2017). The nature of the temperature responses and the acclimation of *R*_M_ are important and much-discussed issues, but they are not considered further here. In contrast, the effects on respiration of woody biomass (the substrate), its accumulation, and the transition rate of respiring sapwood into non-respiring heartwood, have received relatively little attention (Tjoelker *et al.* 1999; Kuptz *et al.* 2011). These latter processes are the focus here.

The fixed-ratio hypothesis of W98 and the scaling hypothesis of R06 could both be used – at least in principle, across the twenty orders of magnitude variation in plant mass – to estimate *R* and *P*_n_ without the need for explicit process-based modelling of *R* (McMurtrie *et al.* 2008; Price *et al.* 2010). However, they may yield quite different results, and both hypotheses (and their supposed underlying mechanisms) have been subject to criticism (e.g. Medlyn *&* Dewar 1999; Mäkelä & Valentine 2001; Kozłowski *&* Konarzewski 2005; O’Connor *et al*. 2007; Keith *et al*. 2010; Agutter *&* Tuszynski 2011; Price *et al.* 2012; Collalti *et al*. 2018, 2019; Collalti *&* Prentice 2019). To our knowledge, there has been no previous attempt to compare these two hypotheses directly, and their consequences for forest carbon balance during stand development, and in the same modelling framework. We attempt to fill this gap by providing illustrative simulations on the long-term trajectories of *R*, *P*_n_ and *P*_n_*:P*, highlighting and discussing the large uncertainty surrounding this issue. The simulations are based on the first principles of mass balance, as adopted in most contemporary vegetation models, and implemented here into a process-based, ecophysiological model that has been tested against detailed time-series observations in an intensively monitored research forest site. We show how alternative assumptions about the live woody turnover (live woody biomass is the metabolically active fraction of sapwood: see Supporting Information) map on to the two alternative hypotheses, while seeking an answer to the pivotal question: is *R* a function of photosynthesis alone (W98’s hypothesis), or of biomass alone (R06’s hypothesis)? Insight into these conflicting hypotheses on plant respiration would help towards a better mechanistic understanding and correct quantification of the stocks and fluxes that determine the carbon balance of forests.

## Materials and methods

### Theoretical framework

A general equation describing autotrophic respiration (*R*) is:

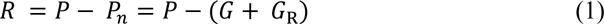

where *P* and *P*_n_ are gross and net primary production, *G* is structural and litter biomass production and *G*_R_ is the flux to NSC reserves and secondary compounds including, exudates and BVOCs (all in g C ground area^−1^ time^−1^). If *R* is further decomposed into growth (*R*_G_) and maintenance (*R*_M_) respiration (McCree 1970, Thornley 1970), then:

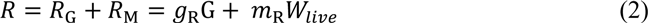

where *g*_R_ and *m*_R_ are the growth and maintenance respiration coefficients (i.e. respiratory CO_2_ released per unit biomass produced by growth and by the maintenance of the existing biomass: both, per unit time and unit mass; Penning de Vries 1975), and *W*_live_ is living biomass (Amthor 2000). *W*_live_ can be broken down further:

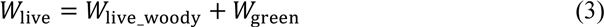

where *W*_live_woody_ and *W*_green_ are the biomass of live woody pools (living cells in stem, branches and coarse roots) and non-woody tissues (leaves and fine roots), respectively. Because plant tissues require N as a component of the enzymes that sustain metabolic processes (including respiration), living biomass is often expressed in nitrogen units, g N ground area^−1^ (Cannell *&* Thornley 2000), while respiration is expressed in carbon units. Then *m*_R_ is in units of g C g N^−1^ time^−1^ (Penning de Vries 1975). Temporal changes in *W*_live_woody_ can be summarized by first-order biochemical kinetics:

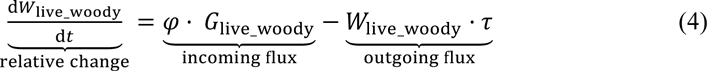

where *G*_live_woody_ is the part of *G* allocated to live woody, *φ* converts carbon to nitrogen content (g N g C^−1^), and *τ* is the live woody turnover rate per unit time (*t*). A similar expression can be written for *W*_green_. The first term on the right-hand side of equation (4) represents the “incoming” flux of new living cells; while second term represents the “outgoing” flux of living cells that die and become metabolically inactive. But while *W*_green_ may be only a small fraction of total forest biomass, not changing much after canopy closure, *W*_live_woody_ (as also total *W*) becomes large during forest development and is potentially a strong driver of *R* (Reich *et al*. 2008). However interpreted and wherever applied, this general approach including a turnover rate parameter (*τ*) is equally valid for any mass-, area- or volume-based analyses (Thornley *&* Cannell 2000).

Setting *τ* = 1 year^−1^ in equation (4) would imply a tight coupling between the previous year’s growth and the current year’s respiration flux – as suggested by Gifford (2003) – and yields a close approximation to the W98 assumption of a fixed ratio between *P*_n_ and *P*, thus cancelling, on an annual scale, any effect of biomass accumulation. The implication of a one-year-lag between carbon fixation and respiration in woody compounds is consistent with the findings of Amthor (2000), Kagawa *et al*. (2006a, b), Gough *et al*. (2008, 2009) and Richardson *et al*. (2013, 2015) of a physiological asynchrony by about one year between *P* and growth (and thus on growth and maintenance respiration).

Alternatively, setting *τ* = 0.1 year^−1^ would imply that most new sapwood cells live for many years, and would closely approximate the R06 assumption of proportionality between *R* and biomass. Thus, the amount of respiring biomass is regulated by the amount of substrate that is produced each year, forming new sapwood, versus the amount that is converted into non-living tissues and no longer involved in metabolism; the balance of these processes being controlled by *τ* (see proofs-of-concept in Fig. 1a and b, and Table 1, for elaboration).

**Fig. 1.**
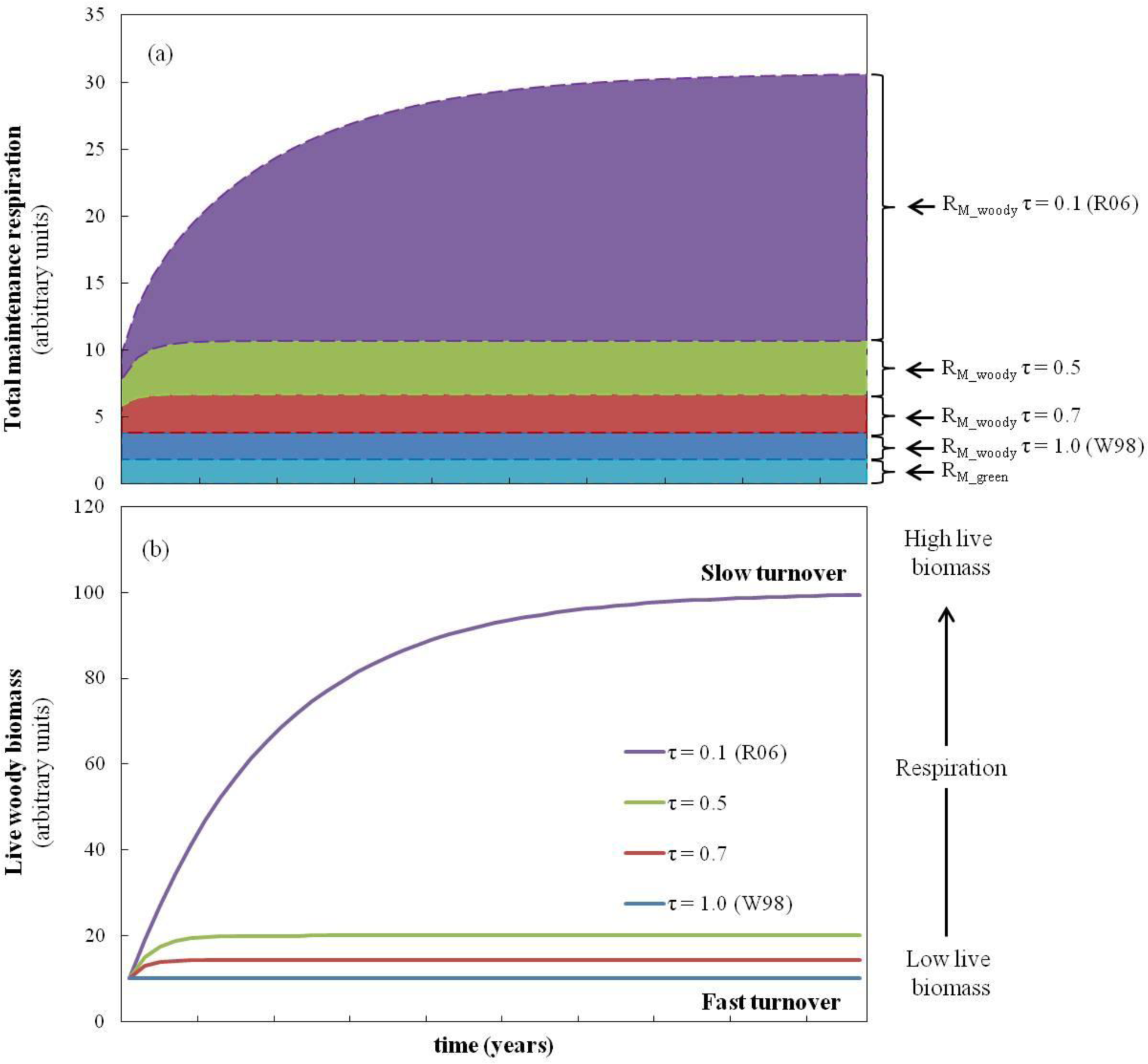
Proofs-of-concept for total *R*_M_ (a) and live woody biomass accumulation (b) over the course of forest development (time) and increases in size, assuming different live wood turnover (τ, yr^−1^) rate values, from 1 (W98) to 0.1 yr^−1^ (R06) and including two intermediate values at 0.5 and 0.7 yr^−1^ (e.g. White *et al*. 2000). *R*_M___green_ (i.e. leaf and fine root *R*_M_) was assumed constant over-time and arbitrarily equal to 2. Summing up *R*_M_wood_ and *R*_M_green_ gives the total *R*_M_. Initial woody biomass was arbitrarily considered equal to 10, new annual live wood was also arbitrarily considered equal to 10, *m*_R_ = 0.2; (*R*_M_ = (*W*_live_woody_ + *W*_green_) · *m*_R_; see Eq. 3). (b) Initial woody biomass was arbitrarily considered equal to 10, new annual live wood was arbitrarily considered equal to 10. The model is: *W*live_woody (t+1) = *W*live_woody (t) + Δ*Win*live_woody (t+1) – Δ*Wout*dead_woody (t+1) (see equation 4 in the main text).

**Table 1.** Underlying modelling assumptions adopted in the analysis

| $\tau$ level | Corresponding underlying assumption | Reference |
| --- | --- | --- |
| $\tau = 0.1$ | Low turnover rate, which implies only accumulation of respiring biomass (*i. e.* $R \propto biomass$) | Reich *et al.* 2006 |
| $\tau = 1$ | High turnover rate, with death of cells annually equalling live cell production (*i. e.* $R \propto P$) | Waring *et al.* 1998 |
| $0.1 < \tau < 1$ | Intermediate turnover rate | e.g. White *et al.* 2000, see Box 2 |
| $\tau = 0$ | Functionally impossible, because it would imply no mortality of cells | - |
| $\tau > 1$ | Physically impossible, because turnover would exceed the number of available living cells | - |

Because carbon supply (photosynthesis) and carbon metabolic demand (respiration) are not necessarily synchronized, the model assumes that temporary carbon imbalances between *P* and *R* (implying *P*_n_ < 0, Roxburgh *et al*. 2005) are met by the remobilization or recycling of NSC stored during previous year(s) – so long as the NSC pool is not completely emptied (the carbon starvation hypothesis; McDowell *et al.* 2008). A full description of the modelled NSC dynamics is provided in Box 1.

#### BOX 1 The function and dynamics of non-structural carbohydrates

NSC is a surprisingly poorly known component of the whole-tree carbon balance, and commonly disregarded in models (Schiestl-Aalto *et al*. 2019; Merganičová *et al*. *under review*). However, the ability of trees to prioritize storage over growth depends on the role of NSC in allowing temporal asynchrony between carbon demand and carbon supply (Fatichi *et al.* 2014). Such imbalances are assumed to be buffered by drawing down NSC reserves. Recent studies support this assumption, showing that during periods of negative carbon balance (for example during the dormant season, periods of stress, or natural or artificially induced defoliation episodes) NSC is remobilized and transported from the sites of phloem loading, while during periods of positive carbon balance plants preferentially allocate recently assimilated carbon to replenish NSC. Only afterwards is “new” carbon used to sustain growth (Weber *et al.* 2018; Huang *et al.* 2019). Because ultimately plant survival depends more on metabolic carbon demands than on growth, some have argued that all positive carbon flows should be used to replenish NSC at the expense of growth until a minimum NSC pool size (30–60% of the seasonal maximum, Martínez-Vilalta *et al*. 2016) is reached (‘active’ storage: Sala *et al.* 2012), thus maintaining a safety margin against the risk of carbon starvation (Wiley *&* Helliker 2012; Huang *et al.* 2019). Note that this assumption departs from the notion that NSC is a mere reservoir for excess supply of carbon relative to growth demand (‘passive’ storage: Kozłowski 1992). In the model, carbon allocation to all tree structural and non-structural pools is computed here daily and is controlled by functional constraints due to direct and lagged C-requirements (Huang *et al*. 2019). It is assumed that a minimum NSC threshold level concentration (11% of sapwood dry mass for deciduous and 5% for evergreen species: Genet *et al*. 2010) has to be maintained for multiple functions including osmoregulation, cell turgor, vascular integrity, tree survival (reviewed in Hartmann *&* Trumbore 2016) and organ-specific phenology (leaf and fine-root formation). The greater the sapwood mass, the greater the minimum NSC threshold must be (Dietze *et al*. 2014). For deciduous trees, four phenological phases are distinguished: (i) the *dormant* phase, where *R* is fuelled by NSC-consumption; (ii) the *leaf onset* phase, when leaf and fine root production consume NSC (unless the carbon balance is positive, in which case new assimilates are used) until the predicted maximum annual LAI is reached; (iii) the *full growing* phase, when new assimilates are allocated to stem, coarse roots, branch, and fruits, and only into the NSC pool if this is below its minimum level; (iv) the *leaf fall* phase, when all assimilates are allocated to the NSC reserve pool while some (∼ 10%) NSC is relocated from falling leaves and dying fine roots (Campioli *et al*. 2013). For evergreen species the model follows a simpler schedule consisting of a first maximum growth phase, when the model allocates NSC to foliage and fine roots up to peak LAI, and a second full growing phase, when the model allocates to all of the pools (Kuptz *et al*. 2011). Such patterns of whole-tree seasonal NSC dynamics have been all recently confirmed by Furze *et al*. (2018) and Fierravanti *et al*. (2019) and a similar phenological and carbon allocation scheme has been adopted by other models (e.g. Krinner *et al*. 2005; Arora *&* Boer 2005).

### Simulation set-up

The logic described above was implemented in a process-based forest growth model (3D-CMCC-CNR), parameterized at site level, and applied, as a case study, to an intensively monitored temperate deciduous forest. Additional model description can be found in Supporting Information, Collalti *et al*. (2014, 2016, 2018; and references therein), and Marconi *et al*. (2017). Very limited data are available on the turnover rate *τ* of live cells in sapwood, which is often either guessed or inferred by model calibration (e.g. White *et al.* 2000). We carried out ten simulations with *τ* varied in arbitrary 0.1 yr^−1^ steps, from *τ* = 1 yr^−1^ (100% of turnover, all the previous year live cells of sapwood becomes non-respiring heartwood in the current year) with *R* mostly depending on the left-hand side term of equation (2) (*R*, down to *τ* = 0.1 yr^−1^ (only 10% of the previous year’s live cells of sapwood biomass dies) and *R* mostly depending on the right-hand side term of equation (2) (*R*. Thus, we started with the largest prior distribution for *τ*, assuming that values outside this range are not functionally possible (Table 1). This approach ensures that any difference in model results reflects difference in specific model assumptions (respiration controlled by photosynthesis or biomass) rather than model structure. We are unaware of any studies reporting changes in *τ* with age or biomass; we have therefore necessarily assumed that *τ* is constant in time.

The standard model configuration assigns *τ* = 0.7 yr^−1^ (Collalti *et al.* 2019) and this same value has been used by several authors in various modelling contexts (e.g. Bond-Lamberty *et al.* 2005; Tatarinov *&* Cenciala 2006). Other models have applied different values (see Box 2). Zaehle *et al*. (2005), Poulter *et al*. (2010) and Pappas *et al*. (2013) found that *τ* is a critical parameter for both LPJ-DGVM and LPJ-GUESS. We are not aware of similar sensitivity analyses for other models. Leaf and fine root turnover rates are assumed here to be 1 yr^−1^, appropriately for deciduous trees (Pietsch *et al*. 2005). The model parameters accounting for ‘age effects’ (e.g. those controlling, among other things, leaf conductance: Kirschbaum, 2000; Smith *et al.* 2001) were set arbitrarily large, to avoid building in prior assumptions. Age- and size-effects are therefore considered synonymous (Mencuccini *et al.* 2005). A stochastic background whole-tree mortality rate (1% of trees removed each year) was retained and included in equation (4) to ensure realistic self-thinning (Smith *et al.* 2001; Kirschbaum, 2005). All other parameters were left unchanged from the standard model configuration.

#### BOX 2 Turnover rates and other uncertainties in models

Most vegetation models assume, among other parameters commonly maintained constant, a fixed rate of sapwood turnover, τ. However, lack of information on this parameter has been already shown to be an important source of uncertainty in the modelled carbon balance of vegetation stands (Goulden *et al.* 2011; Malhi 2012; Collalti *et al.* 2019). Values adopted in current models include: *τ* = 0.7 yr^−1^ in CLM (Oleson *et al.* 2013), Forest v.5.1 (Schwalm *&* Ek 2004), 3D-CMCC-CNR (Collalti *et al*. 2019) and Biome-BGC (Thornton *et al.* 2002); *τ* ∼ 0.75 yr^−1^ in CASTANEA (Dufrêne *et al.* 2005); *τ* = 0.85 yr^−1^ in LPJ-GUESS (Smith *et al.* 2001); *τ* = 0.95 yr^−1^ in SEIB-DGVM (Sato *et al.* 2007), LPJ-DGVM (Sitch *et al.* 2003) and NCAR-LSM (Bonan *et al.* 2003); and *τ* ∼ 1 yr^−1^ in CARAIB (Warnant *et al.* 1994), PnET (Whythers *et al*. 2013), and ORCHIDEE (Krinner *et al*. 2005).

Additional sources of uncertainty include the lack of consideration of a size- or age-related decline in the ratio of living to dead cells (suggesting a declining *τ*) (Damesin *et al.* 2002; Ceschia *et al.* 2002), the effect of changes in climate (which could temporarily increase *τ* to reduce maintenance costs in favour of growth: Doughty *et al.* 2015), changes in tissue N and NSC concentrations (Machado *&* Reich 2006; Thurner *et al.* 2017), and, a probable, genetically controlled down-regulation of basal respiration rates with the ageing of cells (Carey *et al*. 2001; Wiley *et al.* 2017). Moreover, both *τ* and basal respiration rates (*g*_R_ and *m*_R_) are likely to vary among different tree biomass pools (Reich *et al.* 2008). Respiratory carbon losses per unit plant mass may also change to sustain growth as an acclimatory response to carbon demand due to increasing plant size, and perhaps with changing climate (Smith *&* Stitt 2007). These hypotheses are all grounded in theory, but are supported by very limited observations (Friend *et al.* 2014; Thurner *et al.* 2017).

### Test site and model run

The model was applied to simulate 150 years of even-aged stand development in a stand of European beech (*Fagus sylvatica* L.; Sorø, Denmark; Wu *et al.* 2013, Reyer *et al*. 2019) in daily time steps from 1950 to 2100. The reasons for choosing this stand are: (a) the extensive literature on European beech, allowing key parameter values to be assigned with confidence; (b) the exceptional quantity and length of data available at the Sorø site for initializing in 1950 and evaluating the model more than fifty years later, thus, allowing long term processes (including woody biomass accumulation) to emerge; and (c) because the trees are deciduous, we can assume a complete annual turnover of leaves and fine roots and, therefore, more easily disentangle the contributions of *W*_green_ and *W*_live_woody_. Deciduous species are also expected to have greater within-season variability in *P_n_*:*P*, and greater asynchrony in carbon supply and demand, than evergreen species (Dietze *et al*. 2014; Martínez-Vilalta *et al*. 2016). However, both the model assumptions and its results are based on general principles and expected to apply more generally than solely to this specific model and site.

We simulated forest development up to 2100, consistent with the common economic rotation length for this species in northern Europe. After canopy closure, modelled leaf area index (LAI) and the relative amounts of leaf and fine-root biomass became stable or even slightly decrease, as is usually observed (Yang *et al.* 2011, 2016). Therefore, changes in modeled *R*, and its components *R*_M_ and *R*_G_, could be attributed to changes in the total amount of living woody biomass and the costs of its maintenance.

In 1950 the stand was aged 30 years with an average tree diameter at breast height of *∼* 6 cm and a density of 1326 trees ha^−1^. Model state variables were initialized using species-specific functional and allometric relationships from the literature, and previous model applications at this site (Collalti *et al.* 2016, 2018; Marconi *et al.* 2017). Model sensitivity to parameter values and their uncertainties have been assessed in depth in a previous work (see Collalti *et al*. 2019, especially their Fig. 2 and Table 3). Management, in the form of thinning, occurred at the site only up to 2014. After that year, only stochastic mortality was accounted for in the model. Live wood was initialized at 15% of sapwood biomass (as the fraction of current year sapwood: Pietsch *et al.* 2005) and assigned a C:N ratio of 48 g C g N^−1^, not changing with increasing biomass (Ceschia *et al*. 2002; Damesin 2003). The minimum concentration of NSC was assumed to be, ∼11% of sapwood dry mass (Hoch *et al.* 2003; Genet *et al.* 2010; Martínez-Vilalta *et al*. 2016) consistent with measurements on deciduous species (and specifically beech). Daily meteorological forcing variables were obtained as historical ensemble means from five Earth System Models (ESMs) up to 2005 provided by the Inter-Sectoral Impact Model Intercomparison Project (ISI-MIP, Warszawski *et al.* 2014). Data for the period 1995–2005 were then randomly repeated up to 2100. Additional simplifying assumptions were made in order to focus specifically on the effects of increases in tree size, as follows: no disturbances (whether herbivory or management) after 2014; no effect of changes in soil N availability, thus excluding confounding effects of altered N deposition; and, importantly, to avoid possible confounding of temperature effects on *R*_M_ with other warming effects, a stable (1995–2005) climate and atmospheric CO_2_ concentration (*∼* 380 μmol mol^−1^). Exports of carbon to exudates and BVOCs are very slight in this species, and they could therefore be neglected.

**Fig. 2.**
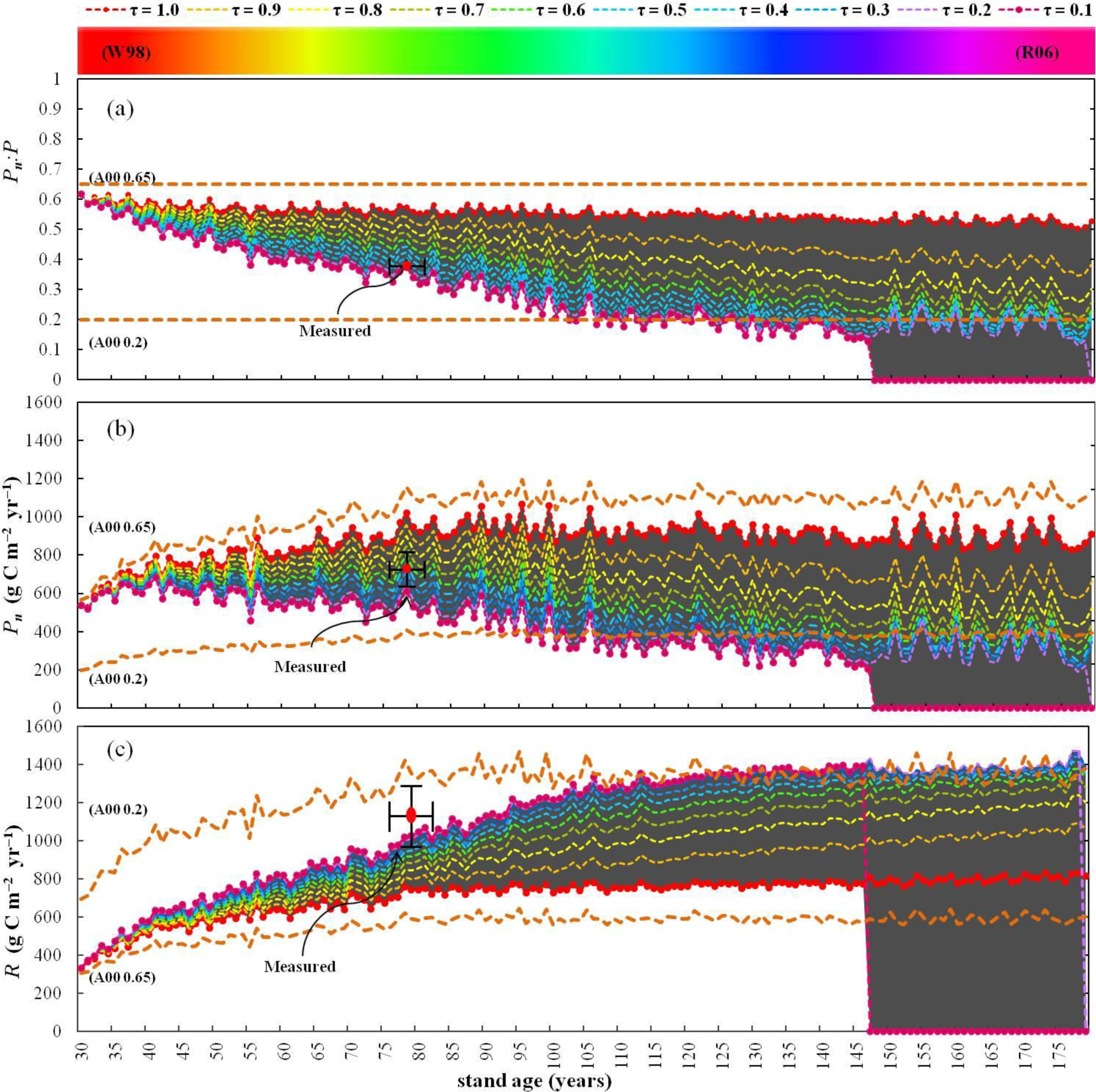
Model results for (a) *P*_n_*:P* ratio (dimensionless), (b) net primary production (*P*_n,_ g C m^−2^ yr^−1^) and, (c) autotrophic respiration (*R*, g C m^−2^ yr^−1^) performed with varying *τ* (coloured lines). The beginning of simulations correspond to 1950 (stand age 30 years); the end of simulations correspond to 2100 (stand age 180 years). The dark-pointed red line can be considered as a mechanistic representation of W98’s fixed *P*_n_:*P* ratio (*τ* = 1 yr^−1^) while the dark pink line approximates R06’s scaling relationship between *R* and biomass (*τ* = 0.1 yr^−1^). Orange dotted lines represent Amthor’s (2000) (A00) ‘*allowable*’ range for the *P*_n_*:P* ratio (0.65 to 0.2). The red dots give the average measured values (Wu *et al.* 2013) at the site for (a) *P*_n_*:P* ratio, (b) *P*_n_ and (c) *R*. Vertical bars represent the standard deviation with horizontal bars representing the period 2006–2010 (stand age ∼ 85–90 years). The shaded area represents the overall uncertainty of model results.

## Results

### Data-model agreement

The standard model configuration satisfactorily reproduced *P*, *R*, *P*_n_ and the ratio *P*_n_:*P* when compared to independent, site-level, carbon balance data (Wu *et al*. 2013) for the period 2006– 2010 (Fig. 2, Table 2), corresponding to a stand age of *∼* 85–90 yrs. *P* was in agreement with eddy covariance data, while *R* was slightly underestimated compared to values in Wu *et al*. Consequently, the model overestimated the average *P*_n_:*P* ratio by 14% compared to Wu *et al*. However, Wu *et al*. argued that the values of *R* they obtained (by subtracting modelled heterotrophic from measured ecosystem respiration) may have been overestimated, given also the large standard deviation (±143 g C m^−2^ yr^−1^). The model results are otherwise in good agreement with Wu *et al*. for woody carbon stocks (both above- and below-ground), annual wood production (the sum of carbon allocated to stems, branches and coarse roots), and annual above- and below-ground litter production (the sum of carbon allocated to leaves and fine roots) (Table 2). Modelled respiration of the woody compartments, leaf and total (above- and below-ground) respiration, and NSC pool and fluxes, are all compatible with values reported by previous investigations, and within the ranges of total, wood and leaf respiration, and *P*_n_:*P* ratios reported for European beech (e.g. Barbaroux *&* Brèda 2002; Barbaroux *et al.* 2002; Knohl *et al.* 2003; Granier *et al.* 2008; Davi *et al.* 2009; Genet *et al.* 2010; Guidolotti *et al.* 2013). A model validation forced by actual measured climate at this site is also described in previous papers (Collalti *et al.* 2016; 2018; Marconi *et al.* 2017).

**Table 2.**
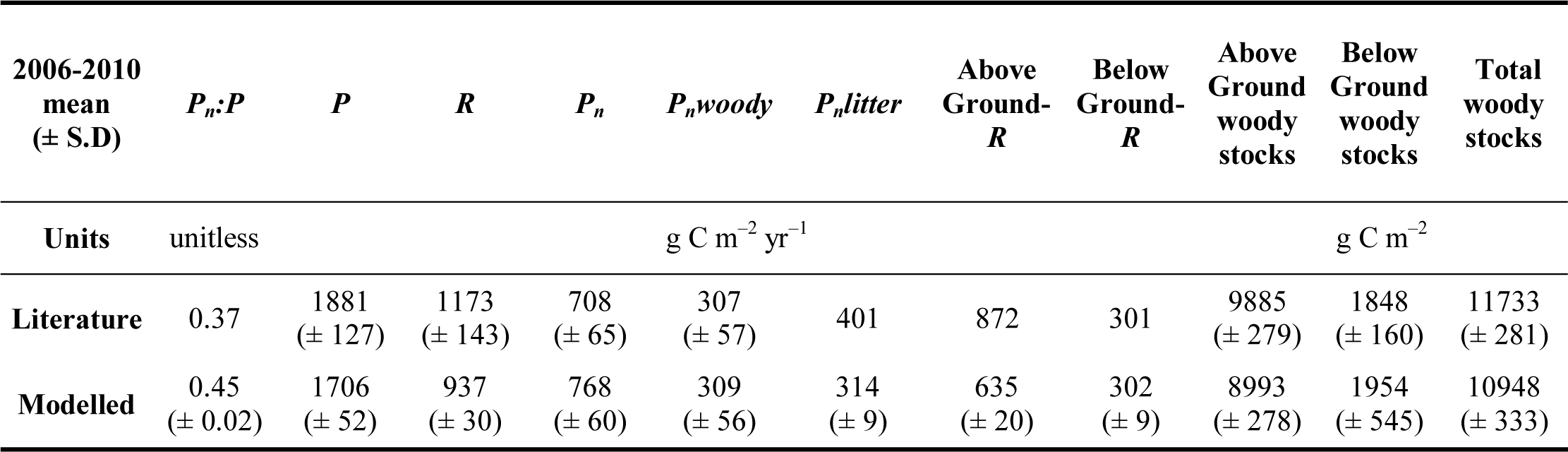
Model validation (averages for the years 2006–2010), in brackets standard deviation (when available). Literature data come from Wu *et al*. (2013).

| 2006-2010 mean<br>(± S.D) | $P_n:P$ | $P$ | $R$ | $P_n$ | $P_nwoody$ | $P_nlitter$ | Above Ground-<br>$R$ | Below Ground-<br>$R$ | Above Ground woody<br>stocks | Below Ground woody<br>stocks | Total woody<br>stocks |
| --- | --- | --- | --- | --- | --- | --- | --- | --- | --- | --- | --- |
| Units | unitless |  |  |  | g C m<sup>-2</sup> yr<sup>-1</sup> |  |  |  | g C m<sup>-2</sup> |  |  |
| Literature | 0.37 | 1881<br>(± 127) | 1173<br>(± 143) | 708<br>(± 65) | 307<br>(± 57) | 401 | 872 | 301 | 9885<br>(± 279) | 1848<br>(± 160) | 11733<br>(± 281) |
| Modelled | 0.45<br>(± 0.02) | 1706<br>(± 52) | 937<br>(± 30) | 768<br>(± 60) | 309<br>(± 56) | 314<br>(± 9) | 635<br>(± 20) | 302<br>(± 9) | 8993<br>(± 278) | 1954<br>(± 545) | 10948<br>(± 333) |

### The effect of varying τ

The simulations produced a spectrum of diverging trajectories, ranging from an approximately steady-state with constant *P*_n_*:P* ratio (for large *τ*) to a constantly decreasing *P*_n_*:P* ratio (for small *τ*) (Fig. 1a). For *τ* = 1 yr^−1^, *P*_n_*:P* stays close to 0.5. For *τ* ≤ 0.2 yr^−1^ *P*_n_:*P* eventually falls below the lower limit of commonly observed values (0.22; Collalti *&* Prentice 2019) and the physiological limit of 0.2 proposed by Amthor (2000). Figure 2 also shows the effects of varying *τ* in determining different trajectories for *P*_n_ (Fig. 2b) and *R* (Fig. 2c) and consequent differences in the partitioning between *R*_M_ and *R*_G_ (Fig. S2) with modelled *R*, at the end of simulations, ranging from ∼ 800 g C m^−2^ yr ^−1^, giving *P*_n_ ∼ 900 g C m^−2^ yr ^−1^ and *P* ∼ 1700 g C m^−2^ yr ^−1^ (Fig. S1 and for NSC flux Fig. S3b) consistent with a steady-state between *R* or *P*_n_ and *P*, to two cases (*τ* = 0.1, 0.2 yr^−1^) in which trees die from starvation.

The model did not generate any consistent power-law relationship between *R* and biomass either for *b* ∼ 1 (i.e. R06), or for ∼¾ (Mori *et al.* 2010), or for ∼⅔ (Makarieva *et al.* 2005), or for ∼⅗ (Michaletz *et al.* 2014) (Table S1). The simulations indicated *b* ∼ 1 initially, shifting with increasing tree size to *b* ∼ 0.74 for *τ* = 0.1 yr^−1^ (*R*^2^ = 0.99, *n* = 117) or 0.19 for *τ* = 1 yr^−1^ (*R*^2^ = 0.84, *n* = 150; ‘*n*’ corresponds to years of simulation). For the relation between *R* and whole-plant N, again the simulations indicated *b* ∼ 1 initially, shifting to *b* ∼ 0.82 for *τ* = 0.1 yr^−1^ (*R*^2^ = 0.99, *n* = 117) or 0.27 for *τ* = 1 yr^−1^ (*R*^2^ = 0.82, *n* = 150) (Figs. 3a and 3c). The highest *b* values corresponded to simulations which ended because the trees died.

**Fig. 3.**
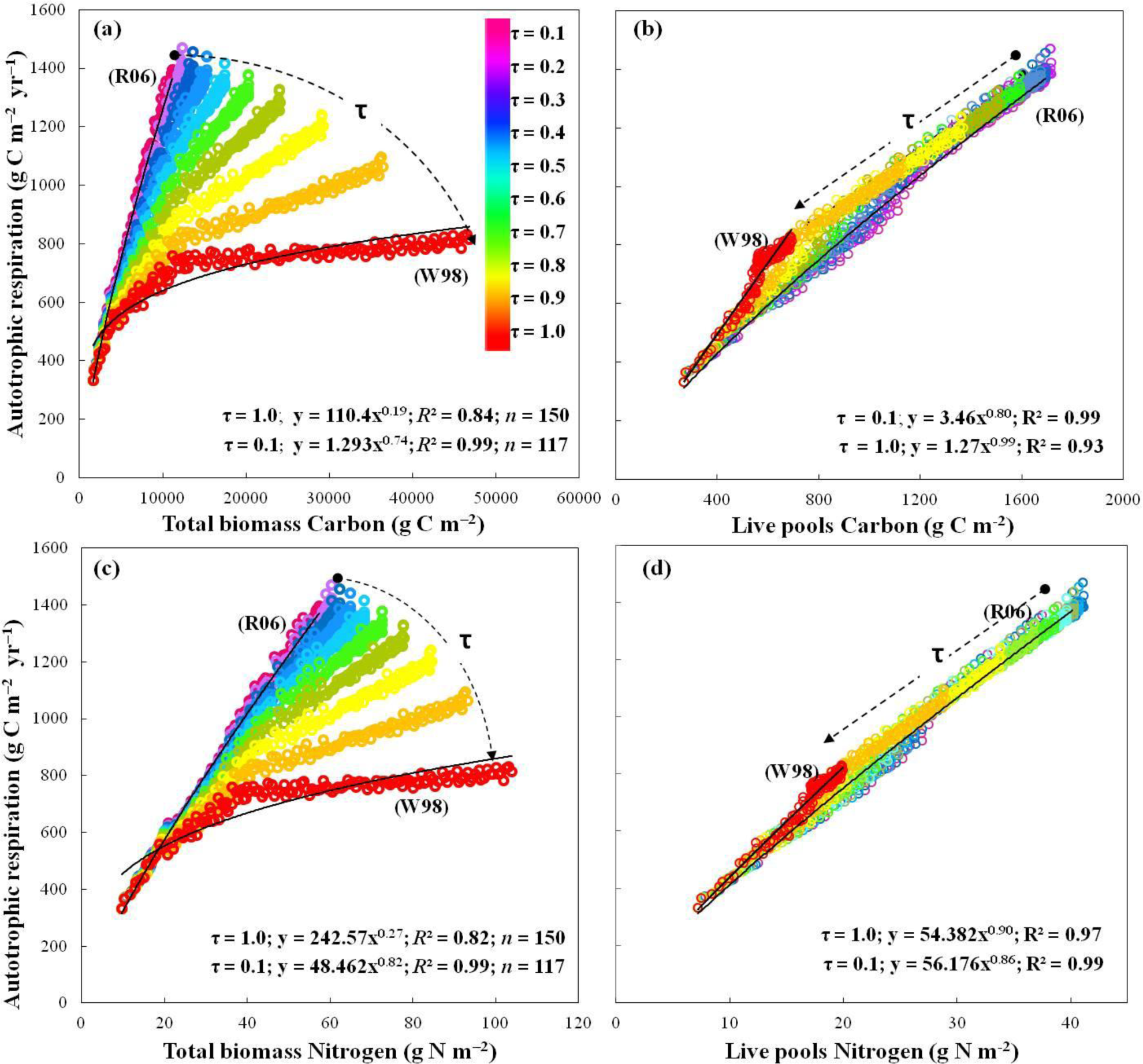
Regression analyses between whole-plant autotrophic respiration (*R*, g C m^−2^ yr^−1^) and (a) whole-plant carbon (*W*; g C m^−2^), (b) carbon in living pools (*W*_live_woody_ + *W*_green_; g C m^−2^), (c) whole-plant nitrogen (*W*; g N m^−2^) and (d) nitrogen in living pools (*W*_live_woody_ + *W*_green_; g N m^−2^). Different colours represent different *τ* values as described in the legend panels (with *τ* = 0.1 yr^−1^, *n* = 117; with *τ* = 0.2 yr^−1^, *n* = 149; otherwise *n* = 150).

## Discussion

### R is not entirely determined by P

A constant *P*_n_*:P* ratio, as implied by W98’s hypothesis and obtained here by setting *τ* = 1 yr^−1^, conflicts with observations from many different tree species that show a substantially slower turnover rate of living cells. In fact, parenchyma cells within secondary xylem are very often more than a year old, and can be up to 200 years old (Spicer & Holbrook 2007). The constant ratio hypothesis is also contrary to the evidence in trees that much of the recently-fixed assimilate pool is at first stored as reserves, and only later used for metabolism or growth (Schiestl-Aalto *et al.* 2015, 2019). Indeed, there are some reports of decoupling between growth (which would imply some CO_2_ released for both *R*_G_ and subsequently *R*_M_) and photosynthesis – with growth ceasing long before photosynthesis – because of the different sensitivities of growth and photosynthesis to environmental drivers. Kagawa *et al*. (2006a) reported for *Larix gmelinii* Mayr. that up to 43%, and according to Gough *et al*. (2009) up to 66%, of annual photosynthetates in bigtooth aspen (*Populus grandidentata* Michx.) and northern red oak (*Quercus rubra* L.) are used during the year(s) after they have been fixed. Gaudinski *et al*. (2009), Malhi (2012), and Delpierre *et al*. (2016) all found negative correlations between annual carbon inflows and above- or below-ground wood growth, from temperate to tropical tree species. Analysing Luyssaert *et al*.’s (2007) global database, Chen *et al*. (2013) found that *R* does not scale isometrically with *P*. Some authors have suggested that *R*_G_ could be supplied exclusively by recent photosynthates while *R*_M_ by previously stored ones (Lӧtscher *et al*. 2004). Along the same lines, Maier *et al*. (2010) for loblolly pine trees (*Pinus taeda* L), Kuptz *et al*. (2011) for beech (*Fagus sylvatica* L.) and Norway spruce (*Picea abies* Karst.), and Lynch *et al*. (2013) for a sweetgum plantation (*Liquidambar styraciflua* L.), found that both *R*_G_ and *R*_M_ are not completely satisfied by recent assimilates, and that some current *R*_M_ can be derived from woody tissues constructed in previous years. Litton *et al*. (2007) and Yang *et al*. (2016) both found low correlations between respiration and the annual production of woody compounds in large datasets. Many studies have also reported little variation in the CO_2_ efflux from sapwood in relation to tree-ring age, despite a stepwise decrease in the fraction of living cells towards the centre of the stems (e.g. Ceschia *et al.* 2002; Spicer *&* Holbrook 2007; Pallardy 2010). These various observations imply that some carbon is fixed one year and used for the tree’s own growth and metabolism in the next or subsequent years, and that the inner sapwood contains a population of living cells formed in previous years.

These observations are all incompatible with the hypothesis of a tight coupling of *R* and *P* (alone), and with model results obtained by assuming complete turnover of live cells in sapwood during a single year.

### R is not entirely determined by biomass

On the other side of the ledger, model simulations indicate that low *τ* values (≤ 0.2 yr^−1^) can lead to excessively high respiration burdens, impossibly low *P*_n_:*P* ratios (< 0.2), and ultimately carbon starvation when all NSC is consumed and whole-tree *R*_M_ or growth can no longer be sustained (Fig. 2). This model result is quantitatively dependent on the values adopted for C:N ratio and the minimum NSC-pool which increases with tree size, but it is consistent with the idea that *P*_n_*:P* ratios ≤ 0.2 are not physiologically sustainable (Amthor, 2000). Amthor described, for a large dataset comprising grasses, tree crops and forest trees worldwide, the 0.65 – 0.2 bounds as reflecting maximum growth with minimum maintenance expenditure (0.65) and minimum growth with maximum physiologically sustainable maintenance costs (0.2). Such a minimum *P*_n_*:P* value agrees also with Keith *et al*.’s (2010) reasoning (analysing *Eucalyptus* forests of south-eastern Australia) that trees always require some annual biomass production in order to survive. With such low *τ*, simulated woody *R*_M_*:R* exceeds 90%, a value much higher than those (56 – 65%) reported by Amthor *&* Baldocchi (2001) (Supporting Information Fig. S2). This situation initiates a spiral of decline, whereby neither *P* nor NSC drawdown are sufficient to avoid a long-term carbon imbalance (Supporting Information, Figs. S3a and b; Wiley *et al.* 2017; Weber *et al.* 2018).

Slightly higher *τ* values (from 0.3 to 0.5 yr^−1^) were found to limit woody biomass increase because of high NSC demand, leading to a shift in the allocation of assimilates to refill NSC at the expense of growth, and *P*_n_:*P* values close to 0.2 (Fig. 2a). Values of *τ* > 0.5 did not show such behaviour and allowed structural and non-structural compounds to accumulate in parallel, while *P*_n_ gradually declined and eventually levelled off. This scenario allows structural biomass accumulation to continue even in older trees, as has been observed (Stephenson *et al*. 2014).

### Scaling relationships

We simulated forest dynamics from juvenile up to very large, mature trees while R06’s results supporting isometric scaling were based on measurements of seedlings and 6- to 25-year-old trees with, presumably, very little heartwood. Some other studies have suggested that the scaling slope of approximately 1 for whole-plant mass may be valid early in stand development, but that the exponent may eventually become smaller than ¾, a phenomenon that has been termed ‘ontogenic drift’ (Makarieva *et al.* 2008). Piao *et al*. (2010) in a global analysis also found a low correlation, and a low scaling exponent, between *R* and whole-plant biomass (*b* = 0.21, corresponding to *τ* ∼ 0.9 in our simulations). Piao *et al*. (2010) argued that, for large mature trees, an increasing fraction of woody C and N biomass is composed of metabolically inactive heartwood, and concluded that a linear-relationship between respiration and whole-plant biomass should not be expected (even if there is a linear relationship of respiration to the live component of woody biomass), while a curvilinear-relationship at the small end of the size-spectrum seemed more appropriate (Kozłowski *&* Konarzewski 2005). Li *et al*. (2005) also found no evidence for an isometric or ¾ power scaling relationship, indicating instead a range between 1.14 and 0.40, decreasing with plant size. The only approximately isometric relationship we found in our simulations – across all *τ* used – was between *R* and the living components of biomass C (*b* in the range of 0.8 to 1, with *R*^2^ always > 0.93) and biomass N (*b* ∼ 0.9, with *R*^2^ always > 0.97) (Figs. 3b and 3d; Makarieva *et al*. 2005; Kerkhoff *&* Enquist 2006; Gruber *et al.* 2009). Conversely, and in accordance with Piao *et al*. (2010), by considering all woody biomass (sapwood and heartwood), *b* consistently deviates from linearity for both C and N in biomass, because – as observed in mature and big trees – an increasing amount of biomass is composed of metabolically inactive tissues that do not respire.

None of these findings are compatible with a tight isometric relationship of *R* to whole-plant C (or N) biomass as proposed by R06.

### R is determined by P, biomass and the demand for reserves

Plants store large amounts of non-structural carbohydrates (potentially enough to rebuild the whole leaf canopy one to more than four times: Hoch *et al*. 2003) and, when needed, plants can actively buffer the asynchronies between carbon demand (i.e. *R* and *G*) and supply (i.e. *P*) by tapping the pool of non-structural carbon (see Fig. S3 in Supporting Information for NSC trends). Several lines of evidence and a growing body of literature support the view of an active sink of NSC. That is, NSC competes with growth, while it controls *R* (and including other non-metabolic functions, see Hartmann *&* Trumbore 2016), in a compensatory mechanism (high NSC demands for respiration means low carbon supply for biomass growth and *vice versa*). Schuur *&* Trumbore (2006) and Carbone *et al*. (2007) for boreal black spruce forest (*Picea mariana* B. S. P), and Lynch *et al*. (2013) for a *Liquidambar styraciflua* plantation, all reported that plant-respired CO_2_ is a mixture of old and new assimilated carbohydrates. Likewise, Vargas *et al*. (2009) for semi-deciduous tree species, Carbone *et al*. (2013) and Richardson *et al*. (2013) for red maple trees (*Acer rubrum* L.), Muhr *et al*. (2013, 2016) for different Amazonian tree species, and Solly *et al*. (2018) for pines (*Pinus sylvestris* L.), beeches (*Fagus sylvatica* L.), spruces (*Picea abies* Karst) and birches (*Betula nana* L.), they all found that old NSC (up to 17 year old) and remobilized from parenchyma cells, can be used for growth or metabolism.

Aubrey *&* Teskey (2018) found that carbon-starved roots and whole-tree saplings die before complete NSC depletion in longleaf pine (*Pinus palustris* L), but the threshold NSC level at which this happens remains unknown for most species. These thresholds are likely to vary among tissues (Weber *et al.* 2018), species (Hoch *et al.* 2003), phenotypes, habit and wood anatomy (Dietze *et al.* 2014), and to increase with tree size (Sala *et al.* 2012). Others have reported that aspen trees (*Populus tremuloides* Michx) cannot draw down NSC to zero because of limitations in carbohydrate remobilization and/or transport (Wiley *et al.* 2017). A minimum NSC level, which has been found to proportionally increase with biomass, may also be required to maintain a safety margin and a proper internal functioning of trees (including osmoregulation), regardless of whether growth is limited by carbon supply (Woodruff *&* Meinzer 2011; Sala *et al*. 2011, 2012; Martínez-Vilalta *et al*. 2016; Huang *et al.* 2019). Genet *et al*. (2010) found for beech and sessile oak (*Quercus petraea* (Matt.) Liebl.) shifts during ontogeny in carbon allocation from biomass growth to reserves regardless of seasonal fluctuations, habitat and climate. Palacio *et al*. (2012) found that black pine trees (*Pinus nigra* Arnold) that were repeatedly defoliated for 11 years, and left to recover for another 6 years, showed reduced growth but similar stem NSC concentration when compared to control trees. Fierravanti *et al*. (2019) found that low NSC accumulation in conifers defoliated by spruce budworm led to a reduction in growth and an increase in mortality.

It has further been suggested that a considerable fraction of NSC (mostly starch) in the inner part of wood may become compartmentalized and sequestered away from sites of phloem loading, and thus no longer accessible for either tissue growth or respiration (Sala *et al.* 2012). Root exudation to mycorrhizal fungi and secondary metabolites (not accounted for here) could also accelerate NSC depletion (Pringle 2016), and potentially create a risk of carbon starvation even for values of *τ* well above 0.2.

Overall, asynchrony between (photosynthetic) source and (utilization) sink implies some degree of uncoupling of *R*, and consequently *P*_n_ (and growth), from *P* and biomass. Carbon demand for metabolism and growth can be mediated by tapping the pool of NSC but only to the extent and to the amount that it is accessible and useable by plants. Therefore, if this active role of NSC can be experimentally confirmed, it will imply that plants prioritize carbon allocation to NSC over growth.

### Implications

It has been suggested that the observed decline of *P*_n_ during stand development cannot be exclusively caused by increasing respiration costs with tree size (Tang *et al.* 2014). The idea, implicit in the growth and maintenance respiration paradigm – that the maintenance of existing biomass (*R*_M_) is a ‘tax’ that must be paid first and which ultimately controls growth – has also been criticized for lack of empirical support (Gifford 2003). While this paradigm has some weaknesses (Thornley 2011), and has not changed much over the last 50 years despite some theoretical and experimental refinements (e.g. accounting for temperature acclimation: Tjoelker *et al*. 1999), it reflects the prevailing assumption embedded in models because, so far, no other general (and similarly promising) mechanistic approach to the modelling of whole-plant respiration has been proposed.

Although plant physiologists are well aware that respiration is neither entirely determined by photosynthesis nor entirely determined by biomass, but rather by plants’ energy requirements for their functioning and growth, we highlight the persistent large uncertainty surrounding this issue in the forestry and forest ecology literature. Both the literature reviewed here and our model results show that any successful modelling approach for plant respiration must necessarily allow plants to steer a middle course between tight coupling to photosynthesis (inconsistent with a carbon steady-state in forest development, and with many observations) and dependence on ever-increasing biomass (risking carbon starvation and death), coupled to the buffering capacity of reserves during carbon imbalances (see Box 1). It seems likely that plants strive to keep an appropriate quantity of living cells that can effectively be sustained by photosynthesis or, when necessary, by drawing on NSC and down regulating allocation to non-photosynthetic, but metabolically active, tissues as to minimize maintenance costs (Makarieva *et al.* 2008). This would suggest active control on carbon use efficiency and on the turnover of the living cells by plants. Yet, despite its importance, NSC use is overlooked in “state-of-the-art” vegetation models. The present study has not been able to provide tight numerical constrains on τ. However, we can unequivocally reject the two, mutually incompatible simplifying hypotheses as both conflict with a large and diverse body of evidence.

Other processes, including hydraulic and nutrient limitations, may be in play (Carey *et al*. 2001; Xu *et al.* 2012). Malhi *et al*. (2015) argued for a link between high whole-plant mortality rates and high forest productivity as ecophysiological strategies that favour rapid growth may also result in fast turnover of trees. However, Spicer *&* Holbrook (2007) noted that metabolic activity does not decline with cell age; and Mencuccini *et al*. (2005) noted that effects of age *per se* (including cellular senescence and apoptosis) are likely not responsible for declining *P*, but are linked to the functional and structural consequences of increasing plant size. This is an important conclusion because it allows models to avoid accounting explicitly for age.

In conclusion, to reduce the large uncertainty surrounding this issue, it will be necessary on the one hand to use models that explicitly account for the turnover of biomass and the reserves usage; and on the other hand, to carry out experimental and field measurements of the dynamics of living cells in wood and the availability of and demand for labile carbon stores. These processes have a direct bearing on the stocks and fluxes that drive the carbon balance of forests.

## Acknowledgements

We gratefully acknowledge Owen K. Atkin, Dennis Baldocchi, Ben Bond-Lamberty, Marcos Fernández-Martínez, Roger Gifford, Francesco Loreto, Maurizio Mencuccini and Richard Waring for reading the many drafts of this paper and having returned many insightful and constructive comments. We thank Andreas Ibrom for providing the site data and for the useful discussions and Carlo Trotta, Angelo Rita, Giulia Mengoli, Elisa Grieco for technical assistance. This work contributes to the AXA Chair Programme in Biosphere and Climate Impacts and the Imperial College initiative on Grand Challenges in Ecosystems and the Environment. It has received funding from the European Research Council (ERC) under the European Union’s Horizon 2020 research and innovation programme (grant agreement No: 787203 REALM). We thank the ISI-MIP project (https://www.isimip.org/) and the COST-Action PROFOUND (FP 1304) for providing the climate historical scenarios and site data used in this work. The 3D-CMCC-CNR-FEM model code is freely available at: https://github.com/3D-CMCC-CNR-FEM

## Author contribution

A.C., M.T., G.H., G.G., G.P., G.M., and I.C.P. designed the study. A.C., G.H., G.G., and M.G.R. carried out modelling work and data analysis. A.C., G.G., M.G.R., G.P., and I.C.P. drafted the manuscript with contributions from all authors.

---

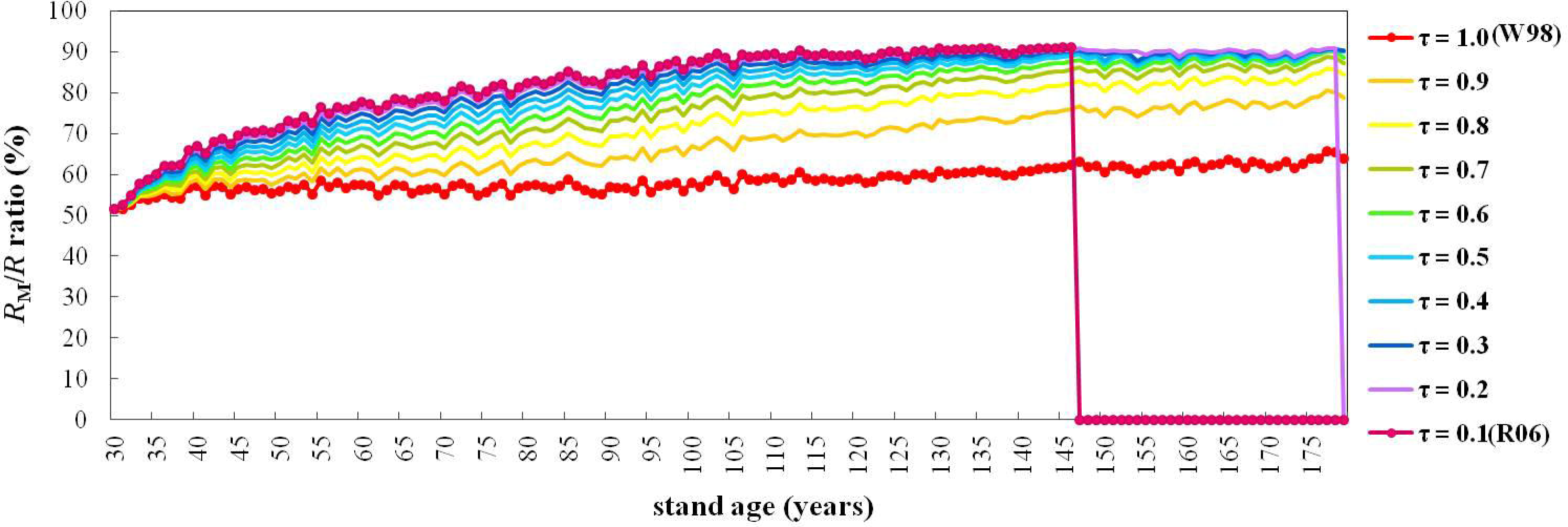
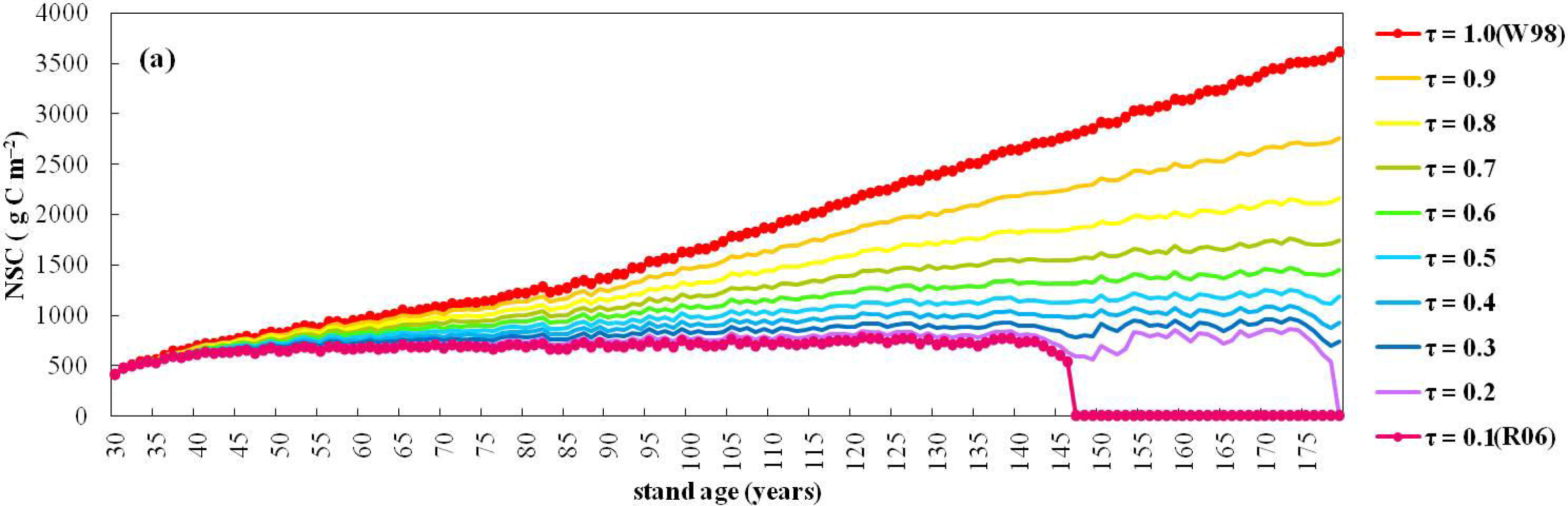
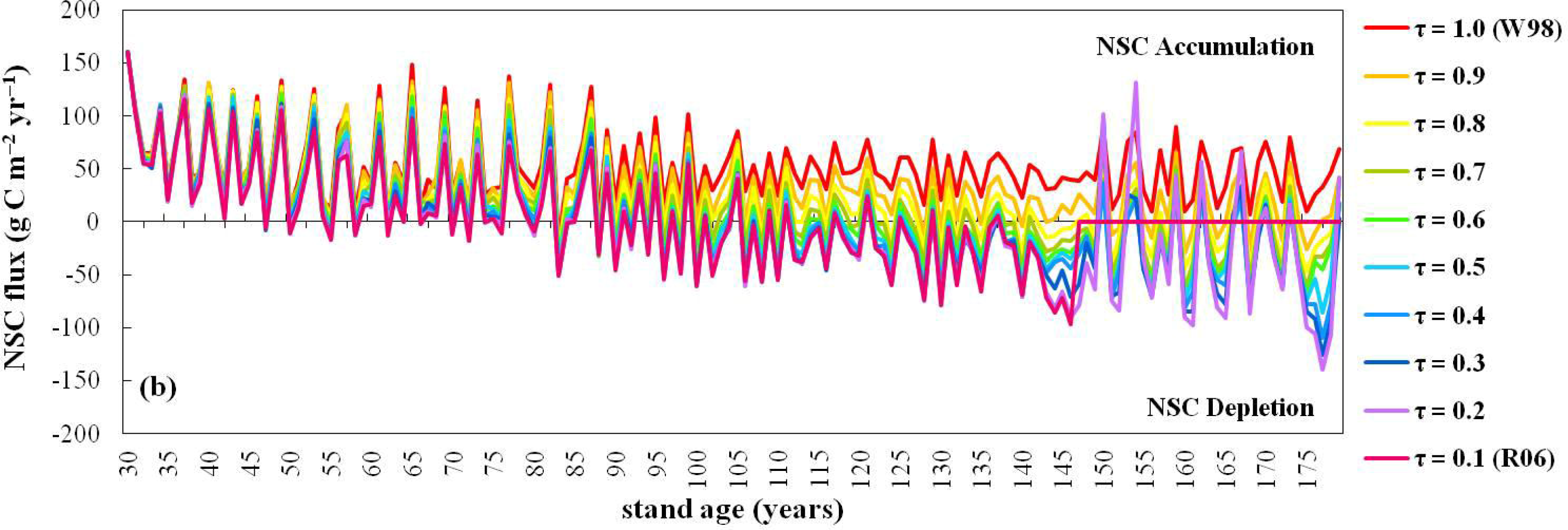

## Notes

#### Summary of Updates

removed one coauthor (erroneously included in the former versions)

