## supporting for "Plant respiration: controlled by photosynthesis or biomass?"

### Supporting Information

#### The model

We applied the first principles described in the main text within a detailed dynamic, processes-based model (3D-CMCC-CNR v.5.5, Three Dimensional – Coupled Model Carbon Cycle, full description is given in Collalti *et al*. 2011, 2014, 2016, 2017, 2018, 2019; Marconi *et al*. 2017). Gross primary production (*P*) is computed using the Farquhar–von Caemmerer–Berry (FvCB) biochemical model (Farquhar *et al*. 1980) as modified for sun and shaded leaves by de Pury *&* Farquhar (1997), with temperature acclimation as in Kattge *&* Knorr (2007). Autotrophic respiration (*R*) is explicitly computed and partitioned into growth (or biosynthesis, *R*_G_) and maintenance (*R*_M_) respiration components (McCree’s 1970) for each carbon (C) pool. Daily net primary production (*P*_n_) is given by *P* minus *R*. Daily structural biomass production (*G*) is given by *P*_n_ minus the C requirement used to fuel *R* and refill NSC reserve pools (*G*_R_). *R*_G_ is considered as a fixed fraction (g_R_, 30%: Larcher 2003; Damesin *et al*. 2002; Damesin 2003; Marthews *et al*. 2012) of new daily assimilates used for structural growth and leaf and fine root production (the allocation and phenological scheme are described in Box 2), while *R*_M_ is computed using an acclimated Q_10_ relationship between ten-day weighted-average temperature (Q_10_ = 3.22 – 0.046*T*; for details on acclimation see Tjoelker *et al*. 2001; Atkin *et al*. 2003; Smith *&* Dukes 2012; Smith *et al*. 2015; Collalti *et al*. 2018) and the nitrogen (N) content of the specific live respiring tissues (the base rate of maintenance is the mass of C respired per unit mass of tissue N content in live mass, *m*_R_ = 0.218 g C g N^–1^ d^–1^; Ryan *et al*. 1991; Drake *et al*. 2011; Oleson *et al*. 2013; Collalti *et al*. 2016, 2018). 3D-CMCC-CNR computes *R*_M_ through a mass-based approach, so increasing live respiring biomass implies an increase in *R*_M_ (Ryan *et al*. 1996; Cox 2001; Ceschia *et al*. 2002).

The main C pools comprise functional and structural compartments: woody biomass (that includes stem, branches and coarse roots, i.e. W_tot_wood_), leaves, fine roots (i.e. W_green_), and the pool of non-structural carbohydrates (NSC) – that is, the reserve pool – which is not accounted for *R*. Among the woody compartments the model distinguishes three main C and N pools (i.e. W_stem_, W_branch_ and W_coarse_root_). Other C and N sub-pools considered are sapwood (W_sapwood_) and heartwood (W_heartwood_) as well as live (W_live_wood_) and dead (W_dead_wood_) woody pools (above- and below-ground parts). W_live_wood_ pool is conceptually different from W_sapwood_ since it includes only live cells, not all of the sapwood biomass – it excludes lignin and cellulose. W_live_wood_ is used to compute respiration, whereas W_sapwood_ is used to compute leaf area index. The relation between leaf area and sapwood follows the pipe model (Shinozaki *et al*. 1964; Mäkelä 1997). Although the model treats W_sapwood_ and NSC pools separately they are interrelated via the minimum NSC content, which is specified as a fraction of sapwood dry mass (Hoch *et al*. 2003; Genet *et al*. 2010; Wiley *et al*. 2017). Newly assimilated C that is partitioned to woody compounds is added to sapwood. New live cells are a fraction of new sapwood, added to previous (and remaining) live cells content after turnover. N allocation follows C allocation according to prescribed C:N ratios (*φ*_pool_), for leaves, fine roots, and woody organs.

As Stockfors *&* Linder (1998) and Ceschia *et al*. (2002) showed that more than 80% of living cells are situated in the outer wood, the amount of new live respiring tissue in the model is a fixed fraction of the sapwood allocated to each pool. Deadwood is the remaining proportion which is not longer involved in metabolic processes. The model relates respiration to the amount of living cellular material rather than assuming respiration to be related to the overall sapwood mass (see for example LPJ-DGVM, Sitch *et al*. 2003; ORCHIDEE, Krinner *et al*. 2005) or to total wood mass or volume (Penning de Vries 1974, 1975; Thornley *&* Cannell 2000; Gifford 2003). The use of the living part to simulate respiration is also adopted by other vegetation models including Biome-BGC (Thornton *et al*. 2002), CASTANEA (Dufrêne *et al*. 2005) and CLM (all versions, Oleson *et al*. 2013).

The live respiring wood biomass is initialized at the beginning of simulation as a fixed fraction of sapwood mass (for stem, coarse root and branch biomass) as in Biome-BGC (Pietsch *et al*. 2005). The quantity of living cells is considered as a fraction of sapwood mass (Hӧlttӓ *&* Kolari 2009; Pallardy 2010). Allometric equations with species-specific parameters convert the initial stem cross-sectional area and sapwood area fraction to sapwood mass (Meinzer *et al*. 2005; Gebauer *et al*. 2008). Juvenile trees stems are thus mostly composed of sapwood mass. The live wood turnover rate parameter *τ* controls the daily rate of live respiring woody biomass that moves from living to dead wood materials the year(s) after its formation and is assumed not to vary over time.

#### References not cited in the main text

Collalti A. (2011). *Sviluppo di un modello ecologico-forestale per foreste a struttura complessa*. PhD Dissertation; University of Tuscia, Viterbo, Italy.

Collalti A., Biondo C., Buttafuoco G., *et al*. (2017). Simulation, calibration and validation protocols for the 3D-CMCC-CNR-FEM: a case study in the Bonis’ watershed (Calabria, Italy). *Forest@ – Journal of Silviculture and Forest Ecology*, 14: 247–256.

Cox P. (2001). Description of the "TRIFFID" Dynamic Global Vegetation Model. Bracknell, Berkshire, Hadley Centre, Met Office, 1–16.

Farquhar G.D., von Caemmerer S., Berry J.A. (1980). A biochemical model of photosynthetic CO_2_ assimilation in leaves of C3 species. *Planta*, doi: 10.1007/BF00386231.

Hӧlttӓ T., Kolari P. (2009). Interpretation of stem CO_2_ efflux measurements. *Tree Physiology*, 29: 1447–1456

Larcher W. (2003). Physiological Plant Ecology. Berlin Heidelberg: Springer-Verlag.

Mäkelä A. (1997). A Carbon Balance Model of Growth and Self-Pruning in Trees Based on Structural Relationships. *Forest Science*, 43(1): 7–24.

Marthews T.R., Malhi Y., *et al*. (2012). Simulating forest productivity along a neotropical elevational transect: temperature variation and carbon use efficiency. *Global Change Biology*, 18: 2882–2989.

Meinzer F.C., Bond B.J., Warren J.M., Woodruff D.R. (2005). Does water transport scale universally with tree size? *Functional Ecology* 19:558–565.

Stockfors J., Linder S. (1998). Effect of nitrogen on the seasonal course of growth and maintenance respiration in stems of Norway spruce trees. *Tree Physiology,* 18: 155–166.

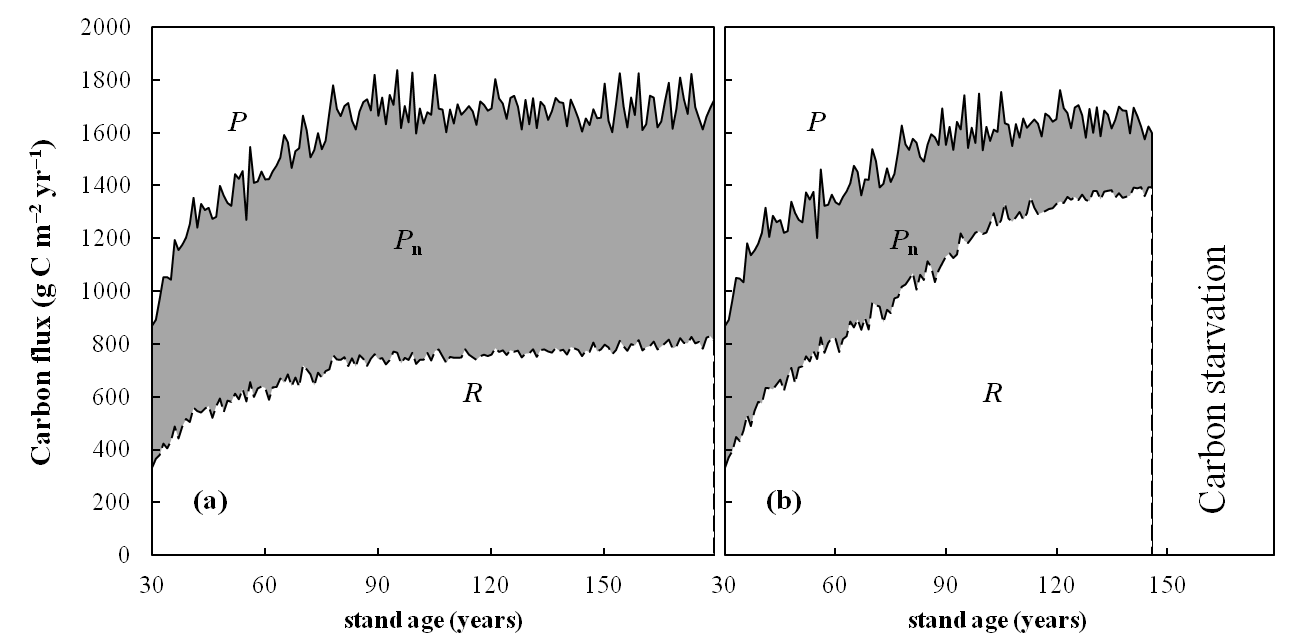

**Fig. S1** The *P*, *P*_n_ and *R* (g C m^–2^ yr^–1^) patterns with stand age at the two bounding extreme turnover rates (*τ* = 1.0 yr^–1^, W98, left panel; *τ* = 0.1 yr^–1^, R06, right panel). Solid lines indicate *P*, dashed lines indicate *R*, the grey area is *P*_n_ calculated as *P* – *R*.

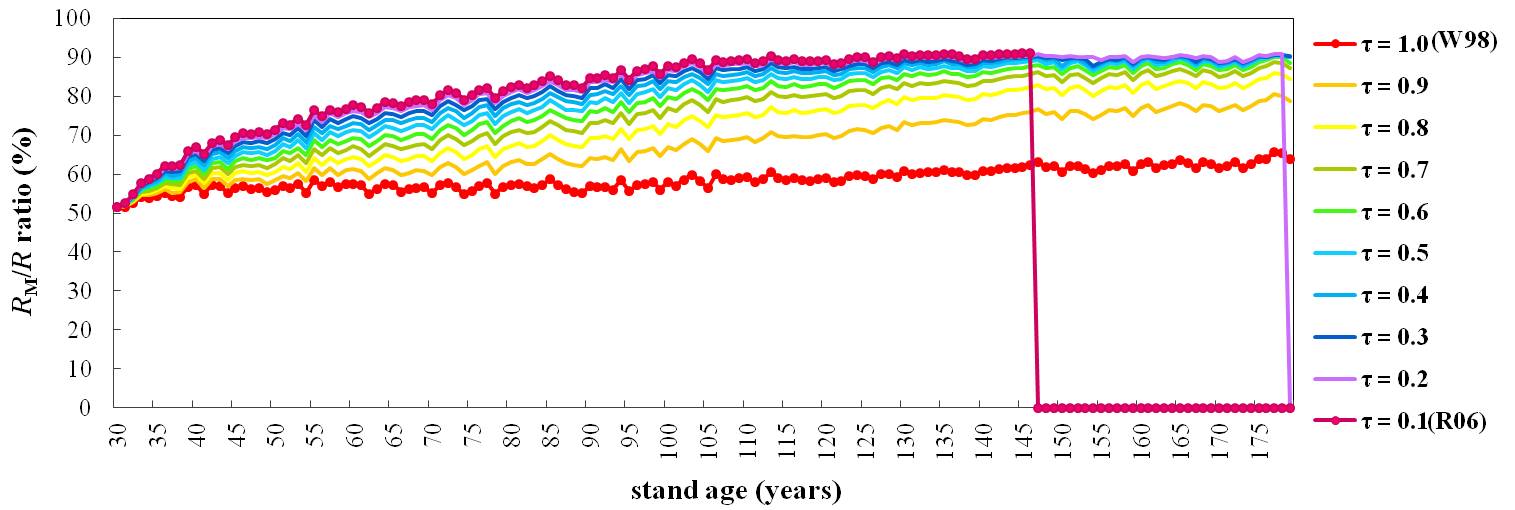

Fig. S2 Model results for annual maintenance (*R*_M_) vs. autotrophic reparation (*R*) ratio (in percentage) (coloured lines) at varying *τ* parameter from 1.0 yr^–1^ (W98) to 0.1 yr^–1^ (R06). The beginning of simulations corresponds to 1950 (stand age 30 years old) while the end of simulations corresponds to 2100 (stand age 180 years old).

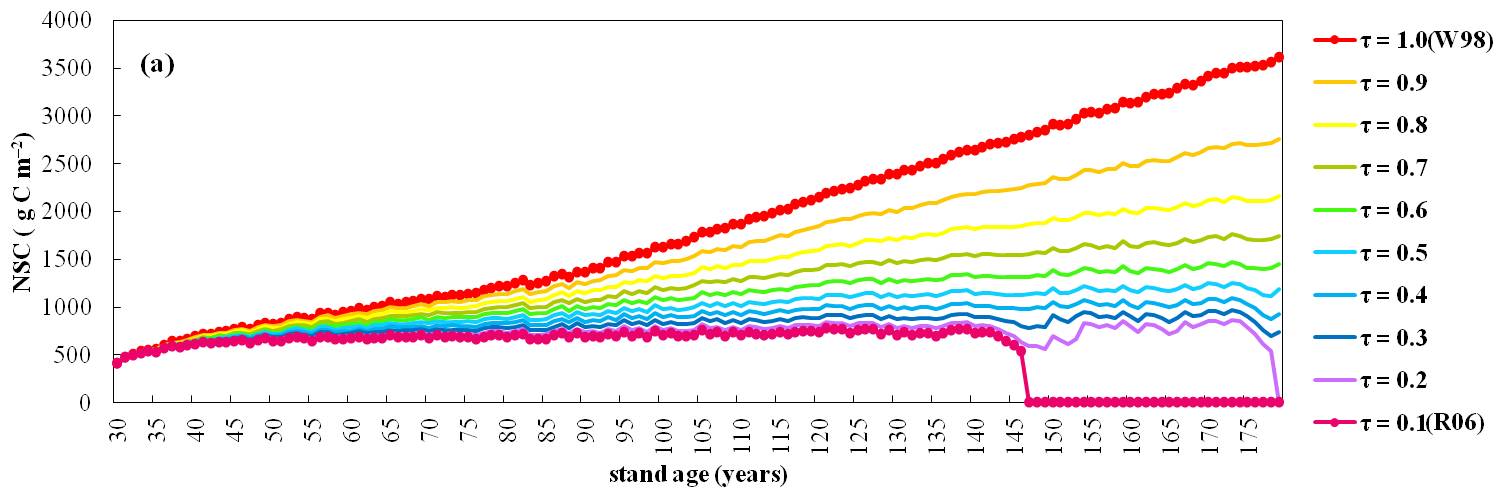

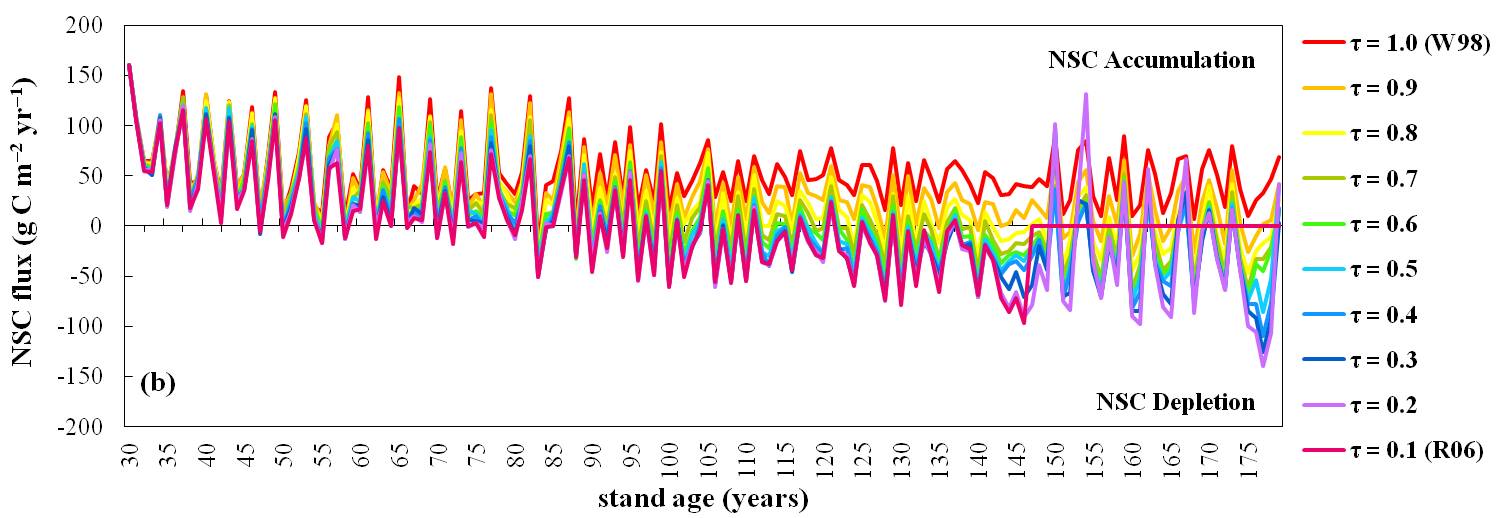

Fig. S3 (a) Non-structural carbon mass (NSC, g C m^–2^ of ground area) and (b) NSC (net) flux (g C m^–2^ of ground area yr^–1^) (coloured lines) trends and interannual variability at varying *τ* parameter from 1.0 yr^–1^ (W98) to 0.1 yr^–1^ (R06). The beginning of simulations corresponds to year 1950 (stand age 30 years old) while the end of simulations corresponds to year 2100 (stand age 180 years old).

Table S1 Scaling exponents (*b*) for annual autotrophic respiration (*R*) against live and whole-plant C and N biomass at varying live wood turnover rates (τ yr^–1^) (with ** *n* = 117, * *n* = 149, otherwise *n* = 150; ‘*n’* represents years of simulation).

|  | *τ* = 0.1 | *τ* = 0.2 | *τ* = 0.3 | *τ* = 0.4 | *τ* = 0.5 | | *τ* = 0.6 | | τ = 0.7 | | τ = 0.8 | | τ = 0.9 | | τ = 1.0 |
| --- | --- | --- | --- | --- | --- | --- | --- | --- | --- | --- | --- | --- | --- | --- | --- |
| Live C | 0.79** | 0.80* | 0.80 | 0.80 | 0.79 | | 0.78 | | 0.77 | | 0.77 | | 0.79 | | 0.99 |
| Total C | 0.74** | 0.72* | 0.68 | 0.63 | 0.59 | | 0.53 | | 0.48 | | 0.40 | | 0.32 | | 0.19 |
| Live N | 0.86** | 0.86* | 0.86 | 0.85 | 0.85 | | 0.84 | | 0.84 | | 0.84 | | 0.84 | | 0.90 |
| Total N | 0.82** | 0.81* | 0.79 | 0.76 | 0.72 | | 0.68 | | 0.63 | | 0.55 | | 0.45 | | 0.27 |
